# Cetylpyridinium chloride-containing mouthwashes reduce the infectivity of SARS-CoV-2 variants *in vitro*

**DOI:** 10.1101/2020.12.21.423779

**Authors:** Jordana Muñoz-Basagoiti, Daniel Perez-Zsolt, Rubén León, Vanessa Blanc, Dàlia Raïch-Regué, Mary Cano-Sarabia, Benjamin Trinité, Edwards Pradenas, Julià Blanco, Joan Gispert, Bonaventura Clotet, Nuria Izquierdo-Useros

## Abstract

Oral mouthwashes decrease the infectivity of several respiratory viruses including SARS-CoV-2. However, the precise agents with antiviral activity present in these oral rinses and their exact mechanism of action remain unknown. Here we show that Cetylpyridinium chloride (CPC), a quaternary ammonium compound present in many oral mouthwashes, reduces SARS-CoV-2 infectivity by inhibiting the viral fusion step with target cells after disrupting the integrity of the viral envelope. We also found that CPC-containing mouth rinses decreased more than a thousand times the infectivity of SARS-CoV-2 *in vitro*, while the corresponding vehicles had no effect. This activity was effective for different SARS-CoV-2 variants, including the B.1.1.7 variant, predominant in UK, also in the presence of sterilized saliva. CPC-containing mouth rinses could therefore represent a cost-effective measure to reduce SARS-CoV-2 infectivity in saliva, aiding to reduce viral transmission from infected individuals regardless of the variants they are infected with.

## BACKGROUND

Several studies have shown the antiviral potential of mouthwashes, which decrease the infectivity of relevant airborne transmitted infectious viruses, such as influenza and distinct coronavirus, including SARS-CoV-2 [1–4]. If proven effective in the oral cavity, this antiviral strategy could represent a globally accessible and affordable measure that could easily be implemented worldwide to reduce the infectivity of SARS-CoV-2 in saliva and to cut the viral transmission chain. Indeed, higher viral loads found in index cases are associated with a significantly higher transmission rates among their contacts [5]. Thus, any intervention aimed at reducing viral loads in the saliva, exhaled as aerosols by infected individuals, could help decrease viral transmission and even prevent superspreading events. As mouthwashes are also available in oral spray formats, they represent a promising strategy for vulnerable populations such as the elderly.

Novel SARS-CoV-2 variants have appeared in different geographical areas, raising concerns about their higher viral transmission potential, the severity of the associated symptoms upon infection, and their ability to escape from pre-established neutralizing responses in individuals already vaccinated [6–9]. In this scenario, the documented broad antiviral efficacy of certain mouthwashes could be instrumental in tackling different SARS-CoV-2 variants and in reducing the impact of their transmission. Yet, and despite the universal applicability of this antiviral approach, and the diverse reports proving the activity of various oral products *in vitro*, we still don’t know which are the individual active components present in these mouthwashes that exert the antiviral effect, and what is their precise mechanism of action. Moreover, we do not know if mouth rinses could be active against different SARS-CoV-2 variants.

Here we focused on the effect of Cetylpyridinium chloride (CPC), a quaternary ammonium compound used in many oral mouthwashes and breath sprays with broad antiseptic and microbicide activity. We compared the anti-SARS-CoV-2 activity of CPC and CPC-containing mouthwashes against their respective vehicles, and found that CPC-containing mouthwashes inhibit SARS-CoV-2 entry into target cells after disrupting the integrity of the viral membrane. CPC-containing mouth washes decreased more than a thousand times the infectivity of replication competent SARS-CoV-2 viruses, were active in the presence of sterilized saliva and effective against different SARS-CoV-2 variants.

## MATERIAL & METHODS

### Cell Cultures

Vero E6 cells (ATCC CRL-1586) were cultured in Dulbecco’s modified Eagle medium, (DMEM) with 10% fetal bovine serum, 100 IU/ml penicillin and 100 μg/ml streptomycin (all from Invitrogen). HEK-293T overexpressing the human ACE2 were kindly provided by Integral Molecular Company and maintained in DMEM (Invitrogen) with 10% fetal bovine serum, 100 IU/ml penicillin and 100 μg/ml streptomycin, and 1 μg/ml of puromycin (all from Invitrogen).

### Pseudovirus production

HIV-1 luciferase reporter pseudoviruses expressing SARS-CoV-2 Spike protein were generated using two plasmids. pNL4-3.Luc.R-.E-was obtained from the NIH AIDS repository. SARS-CoV-2.SctΔ19 was generated (Geneart) from the full protein sequence of SARS-CoV-2 spike with a deletion of the last 19 amino acids in C-terminal, human-codon optimized and inserted into pcDNA3.4-TOPO [10]. Spike plasmid was transfected with X-tremeGENE HP Transfection Reagent (Merck) into HEK-293T cells, and 24 hours later, cells were transfected with pNL4-3.Luc.R-.E-. Supernatants were harvested 48 hours later, filtered with 0.45 µm (Millex Millipore) and stored at -80°C until use. Viruses were titrated in HEK-293T overexpressing human ACE2 to use an equal amount of fusogenic viruses. Pseudoviruses expressing the spike protein containing either the single D614G mutation or the full B.1.1.7 variant were generated and transfected as detailed before [11,12].

### Pseudovirus assay

HEK-293T overexpressing the human ACE2 were used to test mouth rinses and their vehicles at the indicated concentrations. A constant pseudoviral titer was used to pulse cells in the presence of the mouth rinses. After 48h post-inoculation, cells were lysed with the Bright Glo Luciferase Assay system (Promega). Luminescence was measured with an EnSight Multimode Plate Reader (Perkin Elmer). To detect any associated cytotoxic effect, mouth rinse formulations were also mixed with media, and were equally cultured on cells but in the absence of virus. Cytotoxic effects of these products were measured 48h post-inoculation, using the CellTiter-Glo luminescent cell viability assay (Promega).

### Virus isolation, titration and sequencing

SARS-CoV-2 was isolated in march 2020 from a nasopharyngeal swab as described in [13]. The virus was propagated for two passages and a virus stock was prepared collecting the supernatant from Vero E6. Genomic sequence was deposited at GISAID repository (http://gisaid.org) with accession ID EPI_ISL_510689. Compared to Wuhan/Hu-1/2019 strain, this isolate has the following point mutations: 376 D614G (Spike), R682L (Spike), C16X (NSP13) and other 12 in NSP3 (M1376X, P1377X, 377 T1378X, T1379X, I1380X, A1381X, K1382X, N1383X, T1384X, V1385X, K1386X, S1387X). The SARS-CoV-2 B.1.1.7 variant was identified during routine sequencing of a clinical nasopharyngeal swab in Spain in January 2021, deposited at GISAID with accession ID EPI_ISL_900493, and subsequently isolated on Vero E6 cells.

### Viral treatment with CPC-containing mouthwashes and nucleocapsid detection by ELISA

SARS-CoV-2 B.1.1.7 variant and the D614G variant from March 2020 were assayed with Vitis CPC Protect (Dentaid SL), with 2.063 mM of CPC. A total of 250 µl of mouth rinse was mixed with 250 µl viruses for 2 minutes. Untreated viruses were mixed with 250 µl of media for 2 minutes. Mixes with viruses were diluted in PBS and filtered for 10 minutes at 1,000 xg in macrosept^®^ advance centrifugal devices of 100 K MWCO of exclusion (Pall Corporation) to wash away mouth rinses twice. Washed viruses were resuspended in 1.5 ml of fresh media. The amount of SARS-CoV-2 nucleoprotein present in these supernatants was measured with a SARS-CoV-2 Nucleocapsid protein (NP) High-sensitivity Quantitative ELISA (ImmunoDiagnostics), according to the manufacturer’s protocol, but using a 1% BSA buffer instead of the assay buffer of the kit, which contains a detergent to lyse viral membranes and release nucleocapsid content.

### Viral treatment with CPC and dynamic light scattering analysis

100 µl SARS-CoV-2 B.1.1.7 variant or the D614G variant from March 2020 were mixed with 100 µl of CPC (10 mM) or with 100 µl of H_2_O for 2 minutes. These samples were fixed with 1.2 ml of paraformaldehyde 4% (Biotium) for 30 minutes. Particle size distributions of the viral preparations and control vehicles were determined using a dynamic light scattering (DLS) analyzer combined with noninvasive backscatter technology (Malvern Zetasizer; Malvern Instruments, United Kingdom). Samples were measured without dilution. Three different measurements were used to calculate the mean diameter and SD. The effective electric charge on the viral surface was examined by measuring the zeta potential with an electrophoretic mobility and light scattering (ESL) analyzer (Malvern Zetasizer). The samples were placed into the cuvettes and measured without dilution. Three different measurements were used to calculate the mean zeta potential and SD of the dispersed system.

### Antiviral activity

Antiviral activity against SARS-CoV-2 D614G variant from March 2020 was tested against three oral formulations from Dentaid SL containing CPC: Vitis Encias (with 1.47 mM of CPC), Perio Aid Intensive Care (with 1.47 mM of CPC plus 1.33 mM of Chlorhexidine) and Vitis CPC Protect (with 2.063 mM of CPC). Vehicles containing the same formulation but without CPC were also tested in parallel. We also assayed 10 mM of CPC diluted in distilled water. Of note, colorants were removed from all formulations to avoid any interference with luciferase reactions. One ml of mouth rinses or their corresponding vehicles were mixed with 1 ml of SARS-CoV-2 D614G variant for 2 minutes. Virus was also mixed with 1 ml of media as positive control. After 2 minutes of incubation, mixes with viruses were diluted in PBS and filtered for 10 minutes at 1,000 xg in macrosept^®^ advance centrifugal devices of 100 K MWCO of exclusion (Pall Corporation) to wash away mouth rinses twice. Washed viruses were resuspended in 2 ml of fresh media and titrated in triplicates on Vero E6 cells through 10 serial dilutions. After 3 days post infection, cells were assayed in a microscope for viral induced cytopathic effect. To detect any associated cytotoxic effect, mouth rinse formulations were also mixed with media, washed and centrifuged as previously described, and were equally cultured on Vero E6, but in the absence of virus. Cytotoxic effects of these products were measured 3 days after infection, using the CellTiter-Glo luminescent cell viability assay (Promega). Luminescence was measured in a Fluoroskan Ascent FL luminometer (ThermoFisher Scientific). The cytotoxicity obtained for each compound tested determined the limit of detection of the assay.

Antiviral activity against SARS-CoV-2 B.1.1.7 variant along with the D614G variant from March 2020 were assayed with Vitis CPC Protect (with 2.063 mM of CPC) following the same procedure already described. This time, however, 250 µl of mouth rinse was mixed with 250 µl of viruses for 1 or 2 minutes. Untreated viruses were mixed with 250 µl of media and left for 2 minutes. Washed viruses were resuspended in 1.5 ml of fresh media and titrated as previously described. Again, the cytotoxicity obtained for this experiment was used to determine the limit of detection. Finally, antiviral activity against SARS-CoV-2 B.1.1.7 variant in the presence or absence of saliva was assayed with Vitis CPC Protect (with 2.063 mM of CPC). A total of 800 µl of mouth rinse or cell media were mixed with 200 µl of SARS-CoV-2 B.1.1.7 for 30 seconds in the presence or absence of 200 µl sterilized saliva obtained from a non-vaccinated donor with a negative Panbio COVID-19 antigen test (Abbott) 2 days prior saliva donation. This saliva was centrifuged for 20 minutes at 25,000 xg, and then sterilized first with a 0.45 µm and then through a 0.22 µm filter device (Millex Millipore). After 30 seconds of incubation, mixes with viruses were diluted in PBS and filtered for 10 minutes at 1,000 xg in macrosept^®^ advance centrifugal devices of 100 K MWCO of exclusion (Pall Corporation) to wash away mouth rinses twice. Washed viruses were resuspended in 1.5 ml of fresh media and titrated as previously described.

### IC_50_ calculation

Response curves of mouth rinses against pseudoviral entry were adjusted to a non-linear fit regression model, calculated with a four-parameter logistic curve with variable slope. Cells exposed to the pseudovirus in the absence of products were used as positive controls of infection, and were set as 100% of viral fusion to normalize data and calculate the percentage of viral entry inhibition. Cells not exposed to the mouthwashes nor to the pseudovirus were used as positive controls of viability, and were set as 100% to normalize data and calculate the percentage of cytopathic effect. All analyses and figures were generated with the GraphPad Prism v8.0b Software.

## RESULTS

We first tested the capacity of CPC-containing mouth rinses to inhibit SARS-CoV-2 entry into target cells. We employed a luciferase-based assay, using a using a reporter lentivirus pseudotyped with the spike protein of SARS-CoV-2, which allows the detection of viral fusion with target HEK-293T cells expressing human ACE2 receptor. A constant concentration of the reporter pseudovirus containing the SARS-CoV-2 original Spike protein was mixed with increasing concentrations of the indicated CPC-containing mouth rinses, or their corresponding vehicles, and added to the target cells. To control for any mouthwash-induced cytotoxicity, target cells were also cultured with increasing concentrations of the indicated products in the absence of pseudoviruses. By these means, we calculated the concentration at which certain mouth rinses blocked viral entry and achieved a 50% maximal inhibitory capacity (IC_50_). CPC-containing mouth rinses were able to inhibit viral fusion in a dose dependent manner (**Figure 1, A, C, E**, red lines) at concentrations where no cytotoxic effects of the mouth rinses were observed (**Figure 1, A, C, E**, grey lines). Of note, no obvious inhibitory activity was detected on vehicles (**Figure 1, B, D, F**, red lines), clearly pointing to CPC as the antiviral compound contained in the oral formulations. To confirm the specific antiviral activity of CPC, we directly tested this compound resuspended in water, and found that it also inhibited SARS-CoV-2 pseudoviral fusion and entry into target cells (**Figure 1G**). We also assayed the capacity of CPC-containing mouth rinses to reduce the fusion of pseudoviruses displaying different SARS-CoV-2 Spikes, including the D614G mutation and the full B.1.1.7 variant, which has become predominant in UK, and whose higher transmissibility and pathogenicity are a global concern [6]. Yet, a CPC-containing mouthwash was equally effective in abrogating the entry of pseudoviruses expressing each spike these variants (**Figure 1H**). These results indicate that CPC-containing mouth rinses are able to block SARS-CoV-2 viral entry into target cells due to the activity of CPC, which is efficacious against different variants.

**Figure 1.**
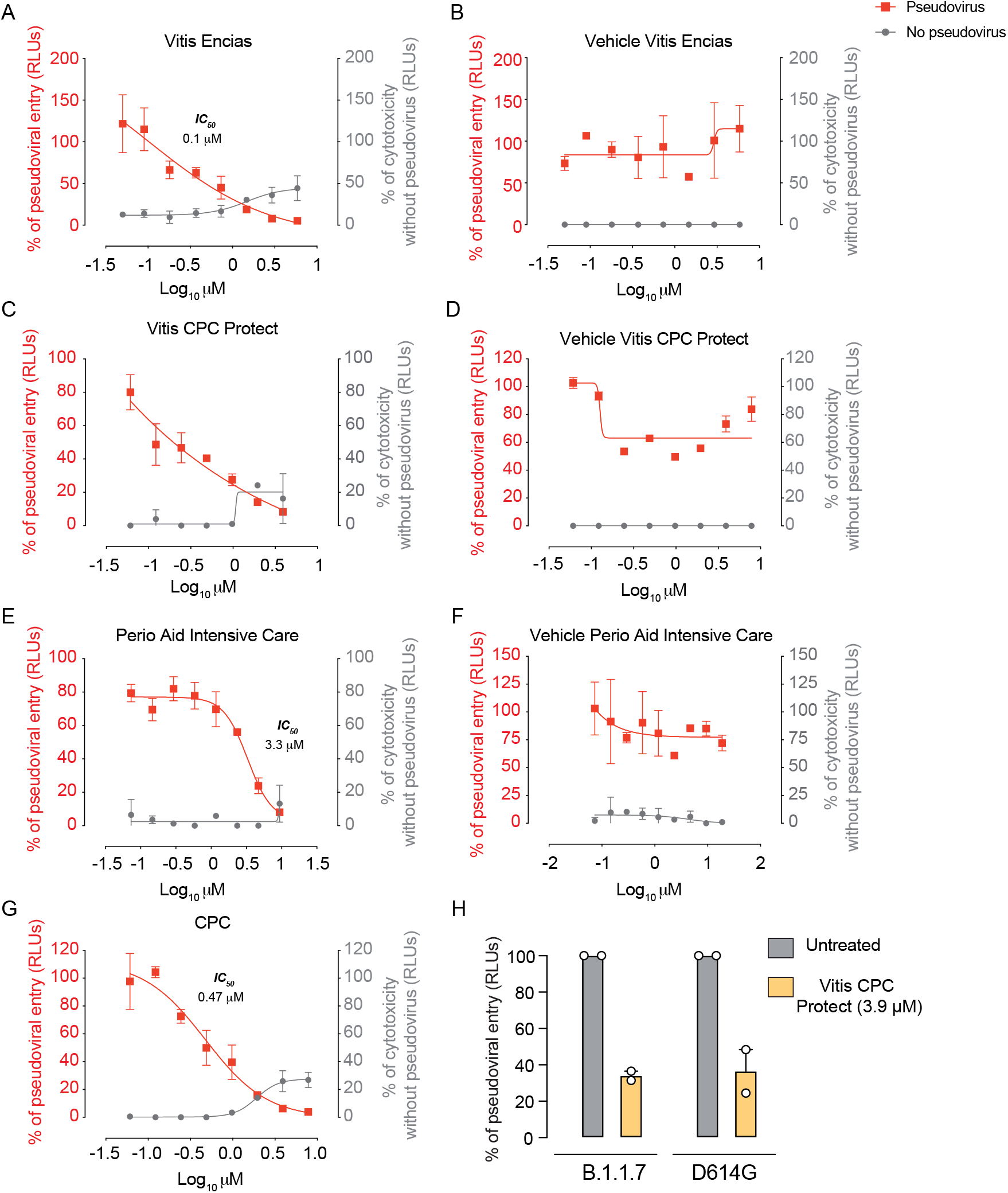
Antiviral activity of CPC-containing mouthwashes inhibiting SARS-CoV-2 entry. Percentage of viral entry inhibition on target HEK-293T cells expressing ACE2 exposed to a fixed concentration of SARS-CoV-2 in the presence of increasing concentrations of oral formulations (**A, C, E**), their vehicles (**B, D, F**) and CPC diluted in water (**G**). Non-linear fit to a variable response curve from one experiment with two replicates is shown (red lines), excluding data from drug concentrations with associated toxicity. When calculated, the particular IC_50_ value of the graph is indicated. Cytotoxic effect on HEK-293T cells expressing ACE2 cells exposed to increasing concentrations of mouthwashes or vehicles in the absence of virus is also shown (grey lines). **H**. Percentage of viral entry inhibition on target HEK-293T cells expressing ACE2 exposed to a fixed concentration of SARS-CoV-2 variants (D614G and B.1.1.7) in the presence of a final concentration of 3,9 µM of Vitis CPC Protect, which had no cell-associated cytotoxicity.

To further understand if CPC-containing mouth rinses abrogated viral fusion by disrupting the viral envelope of SARS-CoV-2, we next worked with two different SARS-CoV-2 clinical isolate variants: the virus circulating in March 2020 in Spain containing the D614G mutation and the B.1.1.7 variant originally detected in UK. Each SARS-CoV-2 variant was mixed at a 1:1 volume ratio with a CPC-containing mouth rinse or left untreated for 2 minutes. To remove the oral rinse, these samples were washed twice with PBS by ultrafiltration using a macrosept^®^ centrifugal device. Collected viruses were assayed with an ELISA that detects SARS-CoV-2 nucleocapsid, but in absence of the detergent used to lyse the viruses in the conventional protocol (**Figure 2A**). In the absence of the lysis buffer, viruses treated with CPC-containing mouthwash were detected to a much higher extent than untreated viruses, regardless of the variant tested (**Figure 2B**). Of note, addition of the ELISA assay buffer containing detergents in untreated viruses increased nucleoprotein detection (**Figure 2C**), but not to the extent of CPC-containing mouthwashes. In addition, we measured the impact of CPC treatment on the zeta potential and size distribution of viral particle s(**Figure 2D** and **Table 1**). Each viral variant was treated with CPC 10 mM at a 1 to 1 volume ratio for 2 minutes, fixed with paraformaldehyde and analyzed by two light scattering methods. Using electrophoretic light scattering (ELS), we found that upon CPC treatment, the zeta potential of untreated viruses that was originally electronegative increased exponentially (**Table 1**). Moreover, this treatment broadened the viral size distribution in both types of SARS-CoV-2 variants (**Figure 2D** and **Table 1**), while CPC or paraformaldehyde alone had a narrower and smaller distinctive profile (**Figure 2D** and **Table 1**), as detected with dynamic light scattering (DLS). The complementary approaches of ELISA and ELS/DLS analyses point out to the effective disruption of viral membranes and loss of electrostatic repulsion, resulting in the emulsion and aggregation of the viral membrane lipids.

**Table 1.** Hydrodynamic size change (Peaks and intensity in %) of different SARS-CoV-2 variants (B.1.1.7 and D614G) and increase in the average zeta potential (mV) of different SARS-CoV-2 variants (B.1.1.7 and D614G) in the presence of CPC obtained by dynamic light scattering. Viruses were mixed with CPC (10 mM) or H_2_O for 2 minutes and fixed with paraformaldehyde. The mean from three different acquisitions was used to measure the mean and SD values represented.

| **Sample** | **Mean particle size** |  |  |  | **Zeta Potential (mV)** |
| --- | --- | --- | --- | --- | --- |
|  | Peak 1 (nm) | Intensity (%) | Peak 2 (nm) | Intensity (%) |  |
| CPC paraformaldehyde | 5,8 ± 1,1 | 56 | 121,2 ± 19,3 | 44 | 15,5 ± 0,9 |
| D614G + H<sub>2</sub>O | 16,1 ± 5,5 | 12 | 226,6 ± 63,7 | 82 | -12,8 ± 1,8 |
| D614G + CPC | 260,4 ± 40,2 | 9 | 1780 ± 369,4 | 91 | 8,9 ± 0,6 |
| B.1.1.7 + H<sub>2</sub>O | 9,2 ± 2,4 | 3 | 237 ± 72,3 | 84 | -11,9 ± 1,4 |
| B.1.1.7 + CPC | 21,2 ± 2,51 | 9 | 1888 ± 231,2 | 91 | 7,87 ± 0,2 |

**Figure 2.**
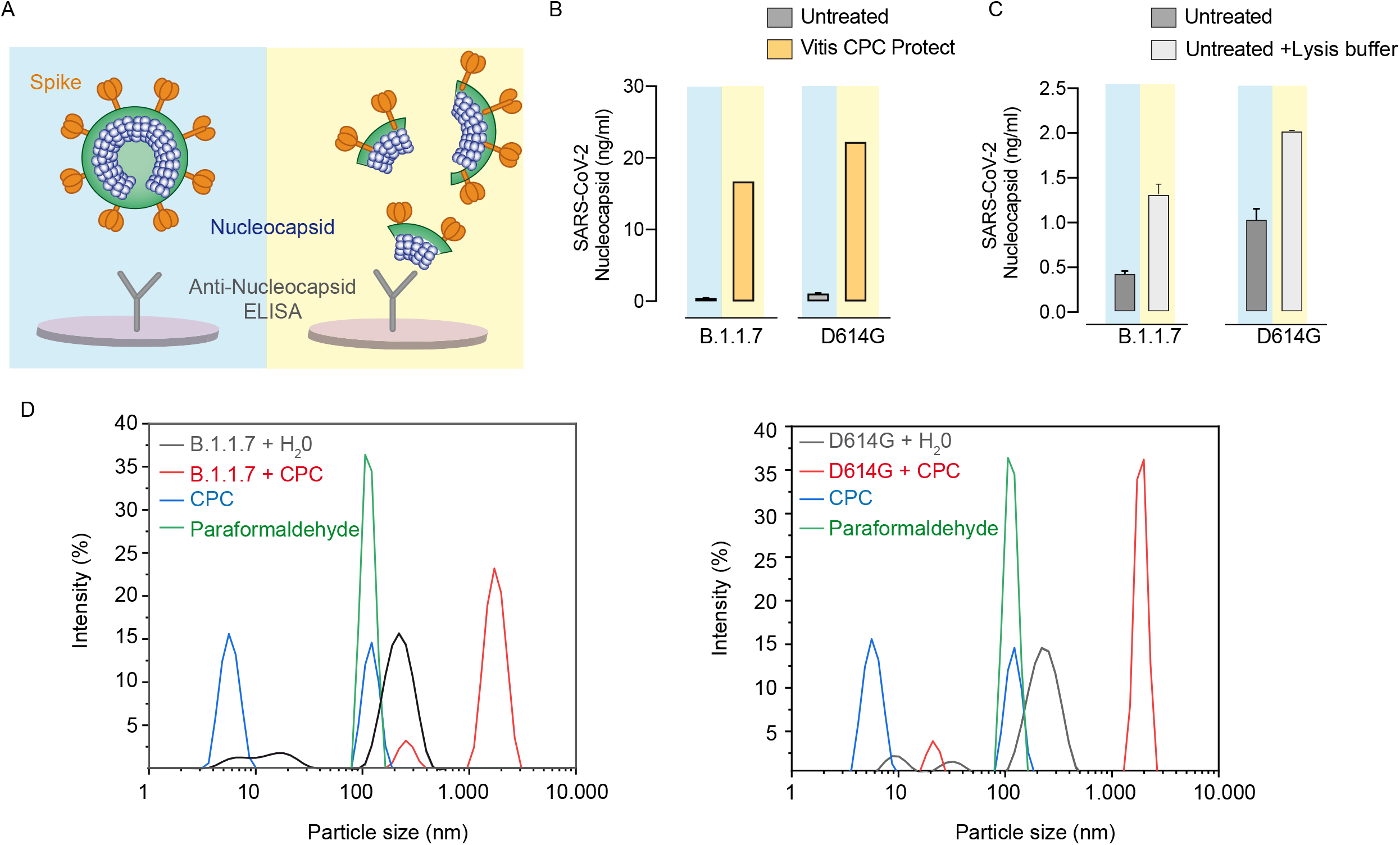
ELISA and DLS analysis of SARS-CoV-2 variants treated with CPC. **A**. Schematic representation of the expected outcome of an ELISA performed in the absence of lysis buffer for untreated viruses (shaded in blue) and those treated with CPC-containing mouth rinse (shaded in yellow) **B**. Amount of nucleocapsid measured after 250 µl of SARS-CoV-2 B.1.1.7 or the D614G variant isolated in March 2020 were left untreated (grey bars) or mixed with 250 µl of CPC-containing mouth rinse (yellow bars) for 2 minutes. Viruses were washed right after treatment by ultrafiltration to remove mouthwashes and were assayed with an ELISA detecting SARS-CoV-2 nucleocapsid performed in the absence of lysis buffer. **C**. Amount of nucleocapsid measured on untreated viruses (grey bars) with the same ELISA in the absence (shaded in blue) or presence (shaded in yellow) of the lysis assay buffer of the kit. **D**. Hydrodynamic size (intensity averaged) of different SARS-CoV-2 variants (B.1.1.7 and D614G) in the presence of CPC obtained by dynamic light scattering. Viruses were mixed at a 1:1 volume ratio with CPC (10 mM) or H_2_O for 2 minutes and fixed with paraformaldehyde.

Next, we tested the capacity of CPC to reduce the infectivity of the clinical isolate of SARS-CoV-2 D614G variant from March 2020. A 1 to 1 volume ratio of SARS-CoV-2 was mixed with CPC, CPC-containing mouth rinses or their vehicles for 2 minutes and washed twice with PBS to remove the formulations by ultrafiltration using a macrosept^®^ centrifugal device. Collected viruses were titrated on Vero E6 cells to calculate the Tissue Culture Infectious Dose 50% (TCID_50_) per ml after each of the treatments. While water used to dilute CPC had no effect on SARS-CoV-2 infectivity (**Figure 3A**), high doses of CPC effectively suppressed viral infection on Vero E6 (**Figure 3A**). Analogously, 2 minutes treatment with CPC-containing mouthwashes decreased about 1,000 times the TCID_50_/ml of SARS-CoV-2, while vehicles had no impact on SARS-CoV-2 infectivity when compared to untreated virus (**Figure 3A**). To control for the presence of mouthwash remaining in the viral preparations that could induce cytotoxic effects, the indicated products were washed in ultrafiltration centrifugal devices, but in the absence of SARS-CoV-2, and equally cultured with target cells for three days. By these means we confirmed that the observed SARS-CoV-2 induced cytopathic effect was effectively inhibited at concentrations where the CPC-containing mouthwashes that could possibly remain after filtration were not toxic for the cells. Similar inhibition was observed when viral stocks were treated with a 10 fold excess volume of CPC-containing mouthwashes for 2 minutes. Thus, CPC exerts an antiviral activity against replicative competent SARS-CoV-2 and CPC-containing mouthwashes have the capacity to reduce 1,000 times the infectivity of a viral stock when treated at least at a 1:1 volume ratio for 2 minutes.

**Figure 3.**
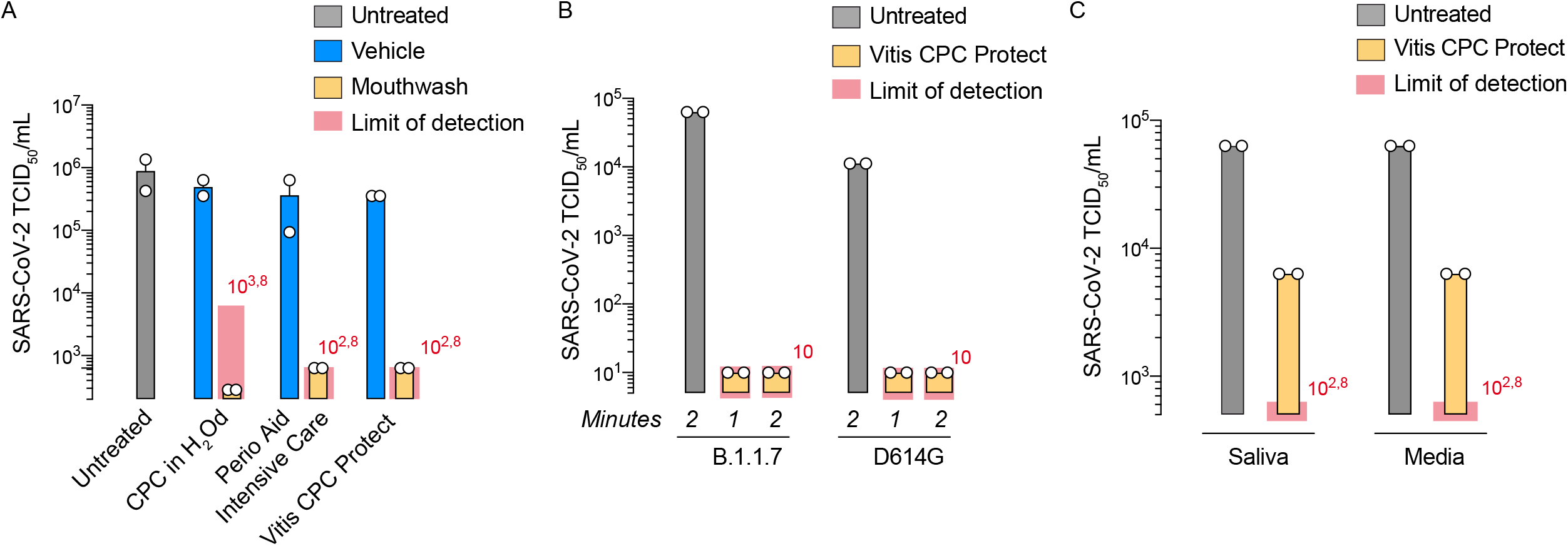
Infectivity of different SARS-CoV-2 variants treated with CPC-containing mouthwashes for different timeframes and with or without sterilized saliva. **A**. One ml of SARS-CoV-2 D614G variant isolated in March 2020 with 10^5.8^ TCID_50_ was treated with CPC (10 mM) or CPC-containing mouth rinses (2 mM) and their respective vehicles for 2 minutes at a 1:1 volume ratio. Untreated virus was used as positive control. Infectivity of treated viruses washed right after treatment by ultrafiltration to remove cytotoxic mouthwashes were assayed on Vero E6 cells 3 days post infection. In parallel, we confirmed that the inhibitory effect was not due to any remaining cytotoxic effect of the mouthwashes, as tested on Vero E6 cells exposed to the media left from washed mouth rinses that were equally centrifuged in the absence of virus. The cytotoxicity obtained for each compound tested determined the detection limit of the assay, which is represented with a shaded red area. Of note, equivalent decrease in TCID_50_ was obtained when viral stocks were mixed at a 1 to 10 volume ratio with CPC-containing mouthwashes (data not shown). **B**. 250 µl of SARS-CoV-2 B.1.1.7 with 10^4.8^ TCID_50_ or 250 µl of SARS-CoV-2 D614G variant isolated in March 2020 was treated with a CPC-containing mouth rinse (2 mM) or left untreated for 1 to 2 minutes at a 1:1 ratio. Infectivity of treated viruses washed right after treatment by ultrafiltration was performed as described in **A**. The cytotoxicity obtained for this mouthwash determined the detection limit of the assay, which is represented with a shaded red area. **C**. 200 µl of SARS-CoV-2 B.1.1.7 was treated with a CPC-containing mouth rinse (2 mM) or left untreated for 30 seconds at a 1 to 10 volume ratio in the presence or absence of 200 µl of sterilized saliva. Infectivity of treated viruses washed right after treatment by ultrafiltration was performed as described in **A**. The cytotoxicity obtained for this mouthwash determined the detection limit of the assay, which is represented with a shaded red area.

To confirm if CPC could reduce the infectivity of different clinical isolates of SARS-CoV-2, we performed and additional experiment including the B.1.1.7 SARS-CoV-2 variant along the D614G circulating variant in March 2020 (**Figure 3B**). In this experiment, we also tested a 1 minute treatment as mouthwashes are usually recommended to be used for 1 minute. After treatment, viruses were titrated as described above, controlling again for the possible presence of mouthwash remaining in the viral preparations that could induce cytotoxic effects. Once again, we could see a reduction of infectivity above 1,000 times regardless of the variant employed or the duration of exposure (**Figure 3B**).

Finally, we tested the capacity of a CPC-containing mouth rinse to exert its activity in the presence of sterilized saliva and using a very restrictive treatment duration of 30 seconds. A 1 to 10 volume ratio of the B.1.1.7 SARS-CoV-2 variant was mixed with CPC-containing mouth rinse or media, in the presence or absence of a 1 volume ratio of sterilized saliva. Viruses were washed twice with PBS to remove the formulations by ultrafiltration, and assayed on Vero E6 as previously described. Treatment for 30 seconds with CPC-containing mouthwashes decreased 10 fold the TCID_50_/ml of the B.1.1.7 SARS-CoV-2 variant as compared to untreated virus (**Figure 3C**). Moreover, the presence of absence of saliva did not alter the inhibition (**Figure 3C**), showing that the CPC-containing mouth rinse has the same antiviral activity in the presence or absence of saliva. Collectively, these results support the potential effectiveness of CPC-containing mouth rinses to decrease viral loads in the oral cavity of infected individuals, regardless of the SARS-CoV-2 variant they are infected with.

## DISCUSSION

Here we have shown that CPC has an antiviral activity against different variants of SARS-CoV-2, and that this compound exerts its activity by blocking viral entry or by inhibiting viral fusion on target cells. CPC acts by disrupting the integrity of the viral envelope, as previously shown for influenza virus [3], which equally affects distinct SARS-CoV-2 variants. Our results indicate that CPC destabilizes the membrane of the different variants, as detected with the ELISA, via electrostatic interactions where the cationic amino groups of CPC cover the negatively charged viral membranes, as detected by the shift in zeta potential. Surfactant CPC activity triggers viral membrane aggregation and colloidal stabilization of solubilized viral membranes that tend to fuse with oppositely charged CPC-bound membranes, increasing size distribution of treated viruses by DLS analysis. This mechanism has therefore the potential to reduce viral infectivity regardless of the variant tested.

Although CPC-containing mouthwashes could protect the oral mucosa from infection, SARS-CoV-2 most likely infect cells via the upper respiratory tract. Thus, further strategies should consider the use of CPC in nasal sprays to fully achieve the prophylactic potential of this approach. Our results point to the utility of CPC-containing oral rinses to decrease viral load in saliva. We have showed that in a very restrictive experiment, where we mixed equal volumes of highly infectious SARS-CoV-2 viral variants with CPC-containing mouthwashes, these treatments reduced more than 1,000 times the TCID_50_ while corresponding vehicles had no impact. Since virucidal activity with a CPC-containing oral rinse was equally effective when saliva was added, this suggest that CPCcontaining mouthwashes will most likely be active in the oral cavity.

CPC-containing mouthwashes could be a cost-effective measure to reduce SARS-CoV-2 infectivity in saliva, aiding to reduce viral transmission from infected individuals. Performing oral washes for 1 to 2 minutes should be enough to effectively decrease the infectivity of viruses in the saliva, especially during the first 2 weeks after infection, when higher viral titers are detected and individuals are more contagious [14]. Several events where numerous people were infected at the same time, which are considered superspreading events, are related to activities where people were either talking, shouting or singing [15,16]. Indeed, viable viruses were isolated from the saliva of COVID-19 infected individuals [17], proving that exhaled saliva micro-droplets and aerosols are infectious. Future work should address if CPC-containing mouth rinses are able to decrease viral load and infectivity of viruses found in the oral cavity of SARS-CoV-2 infected individuals. If proven effective, CPC-containing mouthwashes should be active against those variants that pose a threat to vaccine efficacy, that may increase transmissibility rates, and could even worsen clinical outcome. While prior studies have shown that CPC has an anti-bacterial activity that last for 3 to 5 hours in saliva [18], forthcoming studies should address the duration of the CPC antiviral activity in the oral cavity. This information will be key to effectively validate this approach as a mean to maintain a reduced infectious capacity of SARS-CoV-2 in the saliva with this cost effective intervention.

## AUTHOR CONTRIBUTION

Conceived and designed the experiments: R.L, V.B, J.G, B.C, N.I-U.

Performed experiments and provided pseudoviruses: J.M-B, D.P-Z, D.R-R, M.C-S, B.T, E.P, J.B.

Analyzed and interpreted the data: J.M-B, D.P-Z, R.L, VB, D.R-R, M.C-S, B.T., E.P, J.B, J.G, B.C, N.I-U.

Wrote the paper: J.M-B, D.P-Z, D.R-R, M.C-S, B.T, N.I-U.

## DATA AVAILABILITY

Data is available from corresponding author upon reasonable request.

